# Cryo-ET Analysis Unveils the Minimum Structural Unit and Hierarchical Ultrastructure of Silk Fibers in Silkworm and Spider

**DOI:** 10.1101/2025.03.12.642793

**Authors:** Kai Song, Haonan Zhang, Xueli Zhang, Yan Li, Ping Zhu

**Author notes:** Correspondence (Y.L.) or (P.Z.). ^4^ These authors contributed equally (Kai Song, Haonan Zhang).

## Abstract

Silkworm silk and spider silk are renowned for their exceptional mechanical properties, with the toughness of spider dragline silk exceeding that of silkworm silk and even outperforming high-performance synthetic fibers such as Kevlar and nylon. The mechanical properties of silk fibers are determined by their ultrastructure, including the size, morphology, and arrangement of their fundamental structural units. In this study, we employed the state-of-the-art cryo-electron tomography to investigate the *in situ* ultrastructure of silkworm silk, spider silk, and artificial silk. Our findings reveal that the fundamental structural unit of silkworm silk is a nanofibril approximately 3.6 nm in diameter, connected by numerous bridges, representing the smallest structural unit currently observable. These nanofibrils are aligned parallel to the fiber axis and organized into a herringbone pattern oriented in a specific direction. Multiple layers of this herringbone pattern stack together to form the final micron-scale fiber. In spider silk, a dense structure featuring nanofibrils is clearly visible, with their long axes tightly aligned parallel to the flow direction, leaving no discernible gaps. By contrast, silkworm silk exhibits larger, regionally varying gaps between nanofibrils, while artificial silk lacks the highly ordered arrangement characteristic of natural silks. Our study provides new insights into silk formation and offers valuable principles for designing ultrastrong silk.

## Introduction

Natural silk, primarily derived from silkworms and spider major ampullate gland, exhibit excellent mechanical properties, including high tensile strength and toughness. This gives them not only tensile strength comparable to that of high-performance steel or Kevlar but also exceptional toughness [^1–3^]. For millennia, silk has been prized in textiles for its smooth texture, luster, and strength. More recently, due to its extraordinary mechanical properties, as well as biocompatibility, biodegradability, and eco-friendly manufacturing methods, silk fibers are becoming a candidate material for biomedical engineering [^4–7^], flexible electronic wearables [^8,9^] and storage medium [^10^], and are attracting increasing attention [^11–20^].

The superior mechanical properties of silkworm silk and spider silk are closely tied to their ultrastructure, specifically the morphology, size and hierarchical assembly of the fundamental building block within the fibers. Decades of research have focused on understanding how silk proteins, stored in a soluble state in the silk glands, transform into insoluble solid fibers with distinct mechanical properties. Two primary models, liquid crystal spinning [^21–25^] and micellar spinning [^26^], have sought to explain this process. The liquid crystalline model proposes that spinning dopes in the spider gland and duct adopt a nematic phase, characterized by rod-like structures that are considered aggregates of spherical silk proteins [^21,23^]. In contrast, the micelle model suggests that natural silk fibroin (NSF) forms micelles, approximately 100–200 nm in diameter, in solution due to its amphiphilic primary sequence [^26,27^]. As the NSF concentration increases, these micelles coalesce into globules ranging from 0.8 to 15 µm in diameter, which subsequently align under shear forces to form fibers [^26,28^]. Despite these models, the precise structural units and assembly process of silk proteins in silk glands remain largely elusive. Our recent studies [^29^] have shown a multi-scale controllable self-assembly spinning mechanism based on the natural silk fibroin in silkworm spinning dope, suggesting that the fundamental elements are flexible nanofibrils, approximately 4 nm in diameter, rather than the previously suggested micellar structure [^26,28,30,31^] or rod-like structure formed by aggregation of globular fibroin molecules [^23^]. Within the silk glands, these fibroin nanofibrils initially exhibit random orientation, but intriguingly, they align into a herringbone pattern in the anterior silk gland near the spinneret. This raises questions about whether this ordered structure persists in the final silk fibers or undergoes further transformation during the spinning process. Thus, it is essential to investigate the ultrastructure of silk fibers for a comprehensive understanding of the spinning process.

Investigating the ultrastructure of silk fibers from the atomic to the macroscopic scales remains a major challenge. At the atomic level, previous studies using X-ray diffraction (XRD), Raman spectroscopy, neutron scattering and solid-state nuclear magnetic resonance have established that silk is a semicrystalline polymer. Its crystalline regions are primarily composed of β-sheets formed by polyalanine, contributing to silk’s strength, while the amorphous region, rich in glycine, provide the toughness of the silk fibers [^32–36^]. However, a consensus on the hierarchical structure of silk from the nanoscale to the macroscale is still lacking. Various models based on atomic force microscope (AFM) [^37–42^], scanning electron microscope (SEM) [^43^] and Anderson light localization [^44^] suggest the presence of nanofibrils with diameters ranging from 20 to 220 nm. Nevertheless, techniques like transmission X-ray microscopy [^45^] and ultramicrotomy [^46^] have failed to detect nanofibrils, with some studies indicating silk fibers consist of spherical units, with 10-200 nm in diameter, the size of which varies depending on the type of silk [^26,30,46–48^]. Neutron scattering and small-angle XRD [^35,49^] suggest a long-range organization in silk fibers, although its nature and relationship to the protein sequence remain unclear.

Despite significant progress, many aspects of the supramolecular structure and assembly process of silk remain unresolved. One major challenge is the lack of suitable techniques for studying the native structure of silk fiber *in vivo*. Furthermore, conventional sample preparation methods, such as chemical fixation, dehydration, resin-embedding and heavy metal staining, often introduce artifacts that result in overly condensed structures, obscuring the fundamental units of silk. The intrinsically dense structure of silk further complicates efforts to achieve its high-resolution ultrastructural characterization.

In this study, we employed cryo-focused ion beam (cryo-FIB) milling and state-of-the-art cryo-electron tomography (cryo-ET) to investigate the *in situ* ultrastructure of silk fibers from silkworms, spiders, and artificial silk in their native states. Cryo-FIB is a cutting-edge technique used to prepare high-quality samples for structural and ultrastructural analysis, particularly in the context of cryo-ET. This method preserves the native structure, enables high-resolution imaging by producing ultra-thin sections, and minimizes artifacts typically associated with conventional sample preparation methods. Our study uncovered the morphological and hierarchical structure of fibroin within silk fibers with unprecedented level. The results revealed that the fundamental structural unit of silk is nanofibrils approximately 3.6 nm in diameter, which share the same morphology as those found in the silk gland. These nanofibrils are arranged in parallel along the fiber axis, forming a herringbone pattern. Multiple layers of this herringbone pattern stack to form the final micron-scale fiber, indicating that the herringbone pattern formed near the spinneret remains stable during fiber formation. In spider silk, a more condensed structure adorned with nanofibrils is clearly observed, with their long axes tightly aligned in parallel along the flow direction, leaving no visible gaps. In contrast, silkworm silk exhibits larger, regionally varying gaps between nanofibrils, while artificial silk lacks the highly ordered arrangement found in natural silks. In summary, gaining a deeper understanding of silk’s ultrastructure provides valuable insights into replicating and improving upon the natural spinning processes of silkworms and spiders, guiding the development of polymer-based materials with tunable properties.

## Results

### Structure of fibroin extracted from silk glands of silkworm

Fibroin, a key component of silk, is understood to exist either as a spherical macromolecular complex with a molecular weight of up to 2.4 MDa, composed of a fibroin heavy chain, light chain, and P25 in a molar ratio of 6:6:1 [^50^], or as a rod-like structure formed by the elongation and aggregation of globular fibroin [^23,30,47^]. In this study, natural silk fibroins (NSFs) were extracted from the lumen of posterior silk gland of silkworms. Gel filtration chromatography and 4-16% SDS-PAGE analysis showed that NSFs are a macromolecular complex formed by the fibroin heavy chain, light chain, and P25 subunits (Supplementary Fig. 1a). Interestingly, 3-16% BN-PAGE analysis showed two distinct bands for freshly extracted NSFs, with higher molecular weight bands gradually transforming into lower molecular weight bands upon the addition of amphipol. This transformation displayed a dose-dependent relationship with amphipol concentration (Supplementary Fig. 1b).

For improved sample uniformity, NSFs were treated at a 1:6 molar ratio for 72 hours (Supplementary Fig. 1b) and rapidly frozen via plunge-freezing. Cryo-ET revealed that NSFs exhibit typical fibrous protein characteristics, displaying high flexibility with a diameter of 4.0 ± 0.6 nm and a length of 32.7 ± 16.4 nm (mean ± SD) (Fig. 1, a-c). While the diameters of individual nanofibers were highly uniform, the lengths varied significantly (Fig. 1c). Subtomogram averaging further revealed that fibroin nanofibers appear as a “string of beads” structure (Fig. 1d, Supplementary Fig. 2), and metal ions are essential for maintaining this structure [^29^].

**Figure 1.**
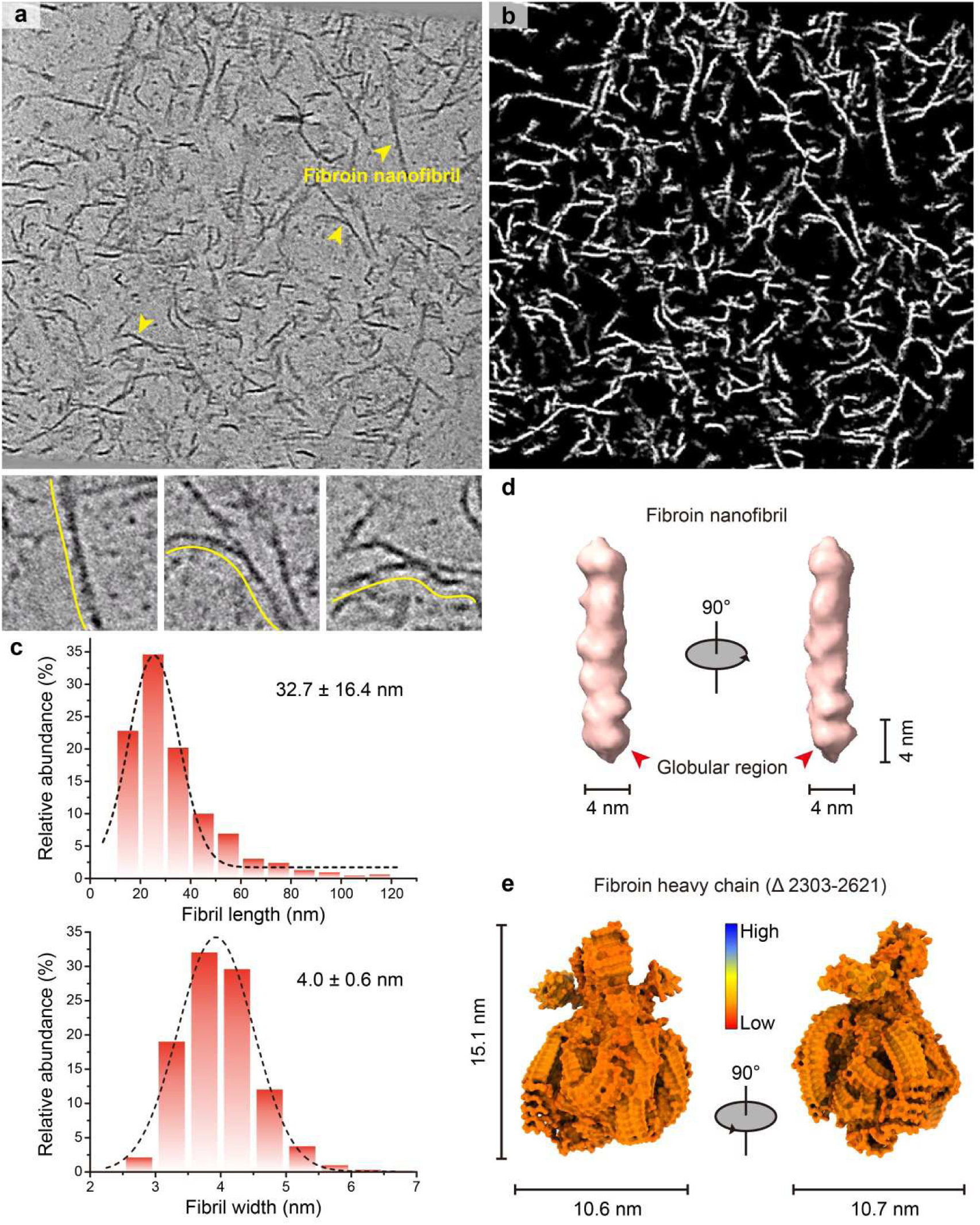
Natural silk fibroin is a beads-on-string structural nanofibril with a diameter of about 4 nm. **a**. Tomographic slice of natural silk fibroin extracted from posterior silk glands. Yellow arrow indicates a typical nanofibril (top), with nanofibrils of varing curvatures shown below (yellow lines depict curved trajectories). **b**. Tomographic slice of segmentation from the same region as in (a). **c**. Nanofibril diameter and length Measurements. (n = 3 tomograms). **d**. The 3D structure of the averaged natural silk fibroin nanofibril in longitudinal view. **e**. Predicted 3D structure of the fibroin heavy chain (Δ2303-2621) by AlphaFold 3.

Most studies suggest that the spherical structures in nanofibrils represent either fibroin monomers [^23,51^] or fibroin aggregates [^26,30,31,47,52–54^]. The fibroin heavy chain consists of 12 repeat sequences and 11 linker sequences, with the repeat sequences containing a high proportion of GAGAGS and GAGAGY motifs that govern the mechanical properties of silk fibers, and the linker sequences showing high conservation (Supplementary Fig. 3), which provides evidence for the concerted evolution of silk protein genes [^55,56^]. Since AlphaFold 3 can only predict polypeptide structures up to 5000 amino acids [^57^], we deleted one repeat and one linker sequence from fibroin heavy chain (Δ2303-2621) to predict its structure. The results indicated that the fibroin heavy chain (Δ2303-2621) can fold into a spherical structure measuring approximately 10.6 nm × 10.7 nm × 15.1 nm (Fig. 1e), which is significantly larger than the 4 nm × 4 nm × 4 nm spherical structures observed in nanofibrils (Fig. 1d). Due to the absence of homologous sequences, the three-dimensional (3D) structure of fibroin heavy chain predicted by AlphaFold 3 lacked accuracy and confidence. Therefore, while this model does not accurately represent the true structure of the fibroin heavy chain, it provides a useful reference for estimating the size of fibroin. That is, if the fibroin heavy chain is folded into a tight spherical structure, its size is approximately 10.6 nm × 10.7 nm × 15.1 nm, while the actual spherical structure in individual nanofibrils is much smaller than predicted size. Therefore, we hypothesize that the smaller globular structures observed in nanofibrils represent individual domains formed by segments of the fibroin molecule. Unfortunately, due to the flexibility of NSF nanofibrils, obtaining high-resolution 3D structures through single-particle reconstruction remains challenging. Consequently, it is still unclear how many spherical domains an individual fibroin nanofibril consist of.

### *In situ* structure of fibroin in silkworm silk

To determine whether the morphology of NSF nanofibrils changes during silk formation, we reconstructed the morphology and organization of silk proteins within silk fibers from 3-molted silkworm. Using cryo-FIB milling, the lamella with the thickness ≤ 100 nm were qualified for decreasing the overlapping signal in the 2D image during the data collection (Supplementary Fig. 4, a-b). The lamella was transferred to a Titan Krios cryo-electron microscope (cryo-EM) for data collection (Supplementary Fig. 4c). In the fibroin region inside the silk fibers, nanofibrils are oriented parallel to the fiber axis, with consistent widths, whereas the sericin region lacked a distinct nanofibril structure (Fig. 2, a-b, Supplementary Fig. 4d). Interestingly, an unknown macromolecular structure was observed in the sericin region (Fig. 2b), but this macromolecular is absent in the sericin region of cocoon silk. It is therefore speculated that this structure might be a specific macromolecule expressed during the early larval stage of silkworms. Despite the overall parallel arrangement of nanofibrils in the fibroin regions, we found that the nanofibril distribution was uneven, with tightly packed areas and looser regions (Fig. 2, a-b). The imperfect parallel arrangement of nanofibrils resulted in gaps within the silk fibers, which could be a contributing factor to fiber fractures during stretching [^58,59^]. In the densely packed areas, nanofibrils were laterally associated, forming bridges (Fig. 2, c-f) that likely contribute to the mechanical properties of silk.

**Figure 2.**
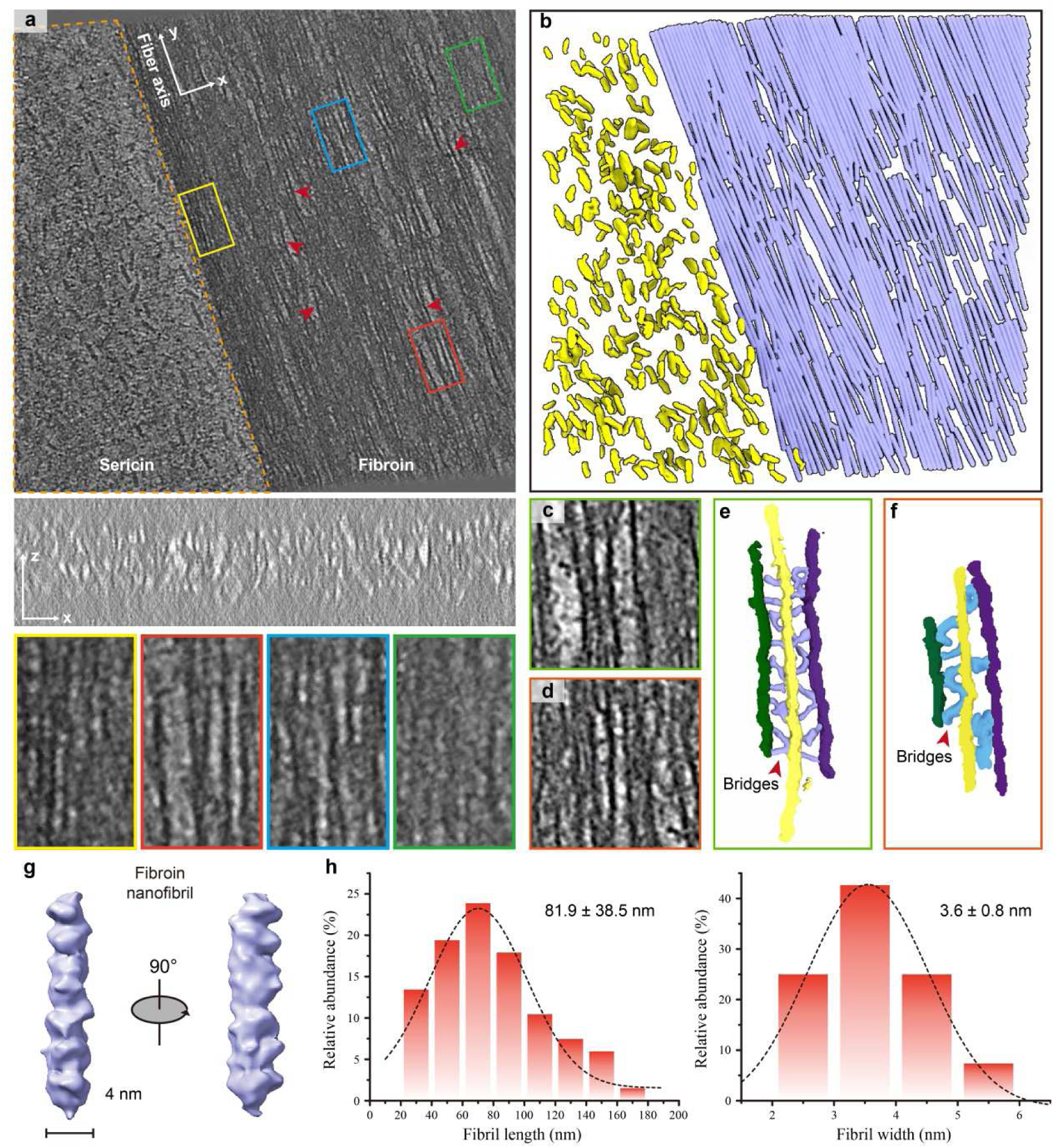
The overall structure of fibroin in silkworm silk remains unchanged. **a**. Tomographic slices showing the morphology and arrangement of fibroin and sericin in the XY and YZ views of silkworm silk. The yellow box: parallel fibroin nanofibrils; red box: non-parallel fibroin nanofibrils; blue box: loosely arranged fibroin nanofibrils; green box: tightly packed fibroin nanofibrils. **b**. The 3D model of fibroin nanofibrils from the same region as in (a). Purple: fibroin nanofibrils; Yellow: Unknow biomolecules. **c-d**. Tomographic slices showing typical cross-bridges between fibroin nanofibrils. **e-f**. 3D segmentation from the same region as in (c-d), illustrating bridges (blue, silver gray) between fibroin nanofibrils (green, yellow, purple). **g**. The 3D structure of the averaged fibroin nanofibrils as the fundamental building block in silks. **h**. Nanofibril diameter and length measurements. (n = 6 tomograms).

A small set of nanofibrils were picked from the tomogram, aligned, and averaged (Supplementary Fig. 5a), We found that the fibroins in silk also appeared as nanofibrils with diameters of 3.6 ± 0.8 nm and lengths of 81.9 ± 38.5 nm (mean ± SD) (Fig. 2, g-h), morphologically similar to those extracted from silk glands *in vitro* (Fig. 1d). The results indicate that the morphology of fibroin nanofibrils remains largely unchanged as they move through the lumen of silk gland to form silk fibers. Our measurements likely overestimate nanofibril length, as tightly connected nanofibrils cannot always be distinguished. Notably, such small diameters for nanofibrils have not been previously reported for silk.

### *In situ* structure of silk protein in spider silk and artificial silk

There are significant differences in mechanical properties between silkworm silk and spider silk, particularly spider dragline silk, which surpasses silkworm silk and even outperforms the best synthetic fibers like Kevlar [^60^]. Using metal shadowing detection, we found that the morphology of silk protein in the lumen of the spider’s major ampullate gland (*Araneus ventricosus*) closely resembled the fibroin of silkworms, appearing as a “string of beads” (Fig. 3, a-c). However, the molecular weight of spider silk protein was lower than that of silkworm fibroin (Fig. 3d). Recent studies have shown that the silk from major ampullate glands of spiders contain 18 proteins [^61^].

**Figure 3.**
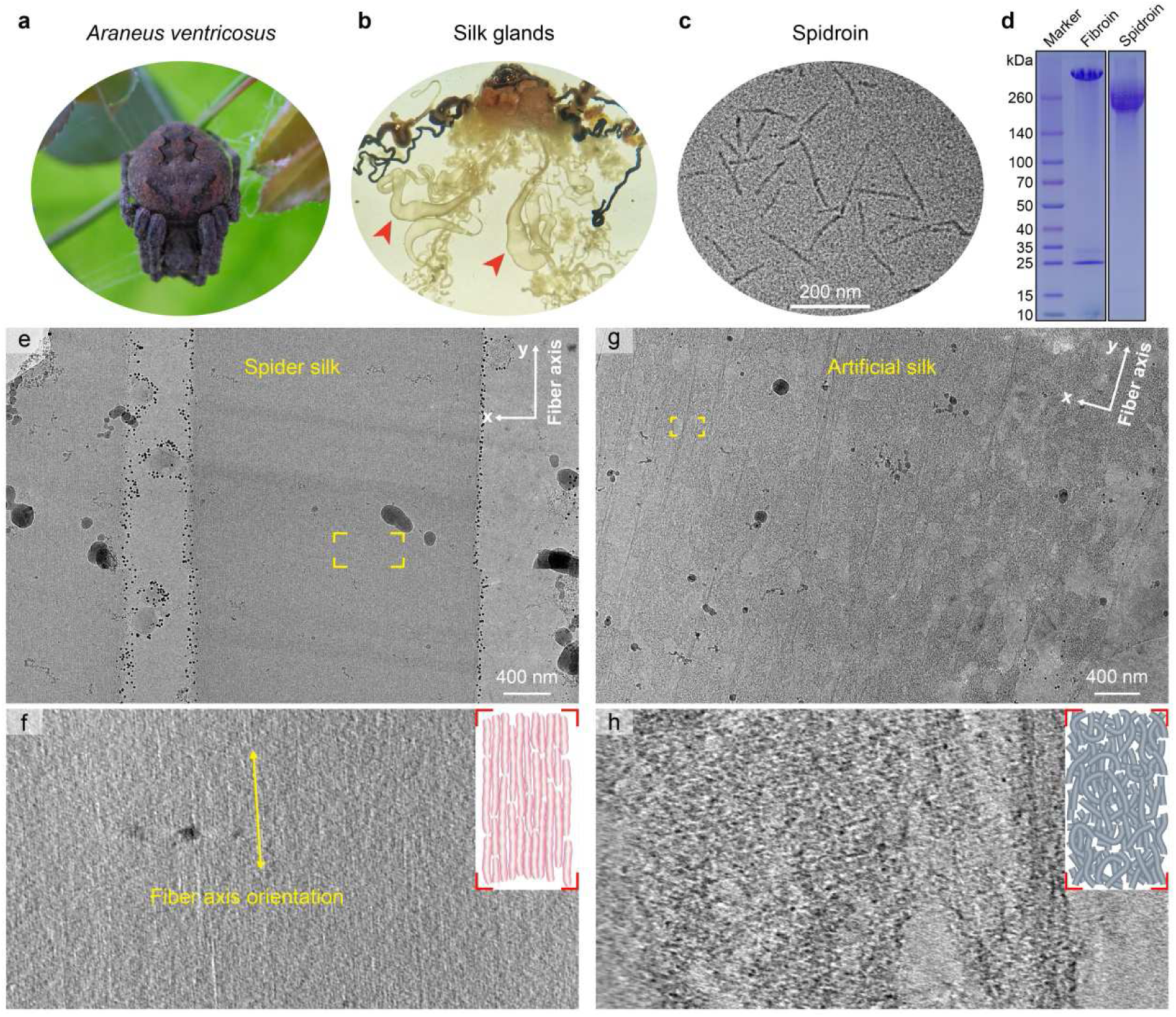
Tightly packed spidroin nanofibrils in spider silk. **a**. Spiders (*Araneus ventricosus*) captured from the wild in Sichuan, China. **b**. Spider silk glands; arrowheads indicate the major ampullate gland. **c**. Metal shadowing of spidroin extracted from the major ampullate gland of spiders, indicating spidroins are nanofibrils. **d**. 4-16% gradient SDS-PAGE analysis of fibroin from silkworm and spidroin from the major ampullate gland of spider. **e**. The low-magnification projection image of densely packed spidroin nanofibrils in spider silks. **f**. Tomographic slice of spidroin nanofibrils enlarged in the position boxed in (e) (not same silk). The inset shows the arrangement of silk protein nanofibrils. **g**. The low-magnification projection image of loosely packed fibroin nanofibrils in aitificial silk. **h**. Tomographic slices of fibroin nanofibrils enlarged in the position boxed in (g). The inset shows the possible arrangement of silk protein nanofibrils.

To further investigate the relationship between the ultrastructure of animal silk and its mechanical properties, we manually pulled fresh silks from the spider and used cryo-FIB milling to prepare lamella ≤ 100 nm thick (Supplementary Fig. 6, a-c). Then, the samples were transferred to a cryo-EM for high-resolution 3D imaging (Fig. 3e, Supplementary Fig. 6d). Cryo-EM revealed numerous nanofibrils arranged parallel to the long axis of the spider silk fibers. Unlike the loose arrangement of nanofibrils in silkworm silk, the nanofibrils in spider silk were tightly packed and well-oriented with no visible gaps (Fig. 3f), resulting in a much denser fiber structure with higher nanofibril density compared to silkworm silk (Supplementary Fig. 7a). This tight packing may explain why spider silk exhibits superior mechanical properties over silkworm silk.

We also prepared artificial silk fibers using the interface wiredrawing method (Supplementary Fig. 8, a-b) and obtained lamella ≤ 100 nm thick via cryo-FIB milling (Supplementary Fig. 8, c-d). In striking contrast, although both natural and artificial silk are composed of fibroin, artificial silk exhibited minimal nanofibril orientation and a non-uniform phase distribution (Fig. 3h, Supplementary Fig. 8e). These findings suggest that the degree of nanofibril orientation and density in silk fibers plays a crucial role in determining their mechanical properties.

### Structural organization of silkworm silk

To investigate whether silk fibers contain ordered structures similar to or higher than the herringbone pattern [^29^] observed near the spinneret in the anterior silk gland, we introduced a deep learning strategy-based method (REST) to establish the relationship between low-quality and high-quality density, applying this knowledge to restore signals in cryo-ET (Supplementary Fig. 5b) [^62^]. The results show that this method improves the signal-to-noise ratio of tomographic slices and reduces some artifacts caused by the excessive density of silks and the missing data region in the 3D reconstruction of sample (missing wedge) (Fig. 4, a-b). Notably, we observed numerous regularly arranged dot-like structures in the XZ view of the silk fibers (Fig. 4, a-b, bottow). By tracking each dot along the y axis, we determined that each dot corresponds to a nanofibril, which were densely stacked on the YZ view (Fig. 4, c-d). This arrangement is consistent with the herringbone patterns found near the spinneret in the anterior silk gland (Fig. 4e) [^29^]. Previous studies [^29,63^] identified an axis within the herringbone pattern, possibly a pseudo-axis formed by interactions involving the N-terminal domain of the fibroin heavy chain of fibroin. In our experiments, we observed both tightly connected regions and blank regions within the same herringbone pattern. Interestingly, a distinct axis perpendicular to the nanofibrils was also evident in the blank regions of the herringbone pattern (Supplementary Fig. 9a).

**Figure 4.**
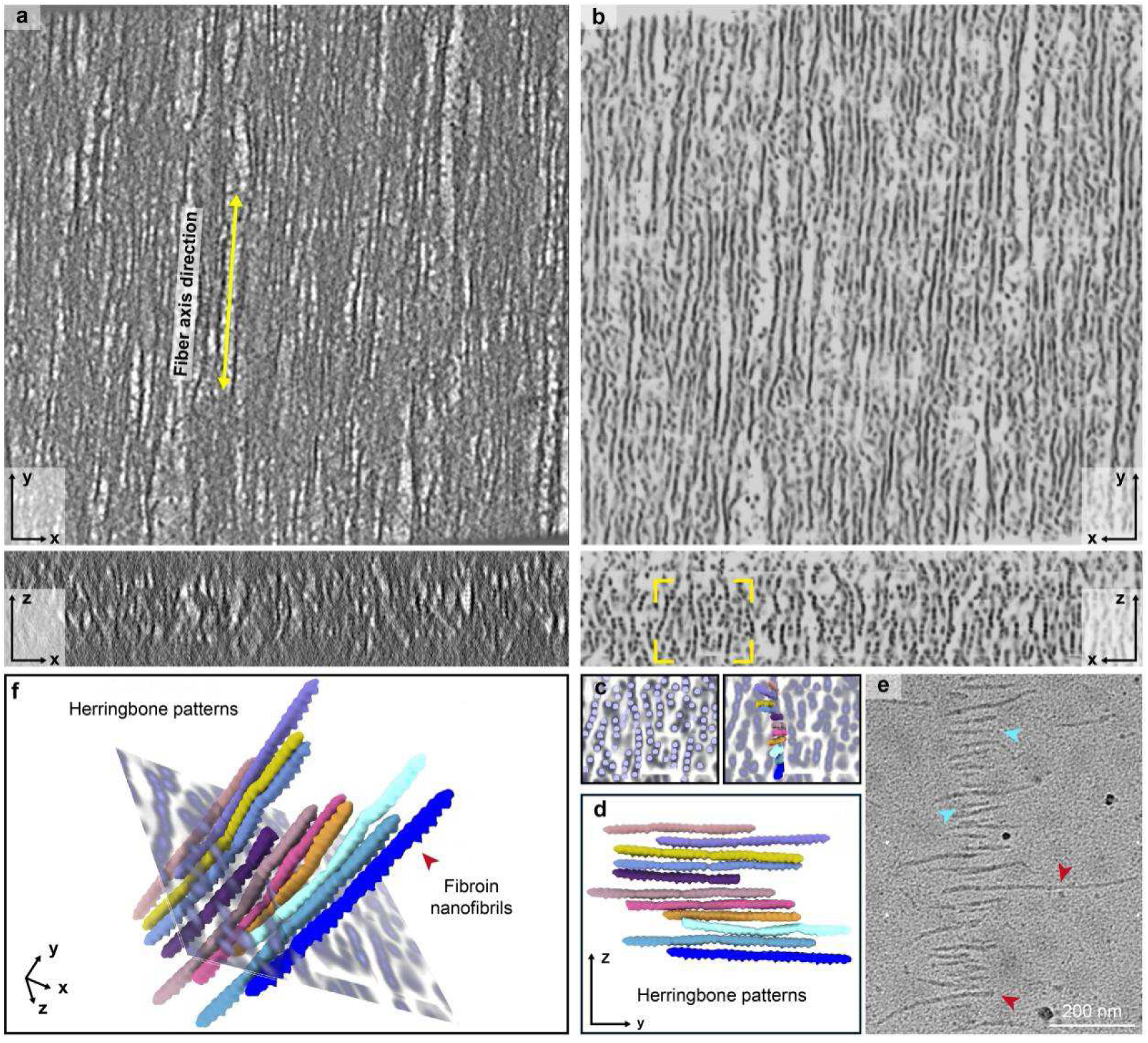
Fibroin nanofibrils form an anisotropic herringbone patterns in silk. **a**. Tomographic slices of fibroin in XY and YZ views of silkworm silk. **b**. Tomographic slices from the same region as in (a) using the REST method, which significantly removes noise and improves signal-to-noise ratio of fibroin nanofibrils, particularly in the XZ view, where point-like structure along the z-axis are evident. **c**. A 3D model of fibroin nanofibril organization in the same slab as in (b), showing a herringbone-like pattern. **d**. Tilted view of the 3D model in (c). **e**. Natural silk fibroin self-assembled into herringbone patterns at the distal region of anterior silk gland near the spinneret. f. Cryo-ET-based 3D model of the herringbone pattern, showing parallel nanofibrils along the z-axis.

Recent studies propose that the herringbone patterns of fibroin may actually resemble a 3D bottlebrush structure [^63^]. Statistical analyses of the distances between points along the z-axis and x-axes revealed that the spacing between points along the z-axis is significantly smaller and more uniform than that along the x-axis (Supplementary Fig. 7b). This finding indicates that the nanofibrils within the silk are anisotropically arranged rather than simply stacked in parallel. Such an arrangement contributes to enhancing the mechanical properties of the material [^64^]. Moreover, they are organized into a herringbone pattern, where multiple layers of this herringbone pattern are stacked along the x-axis to ultimately form micro-sized silk fibers (Fig. 4f). Obviously, the final microscale fiber does not exist as a compressed bundle of aligned nanofibrils, and no higher-order organization was observed beyond the herringbone pattern. This finding suggests that the herringbone pattern, formed near the spinneret in the anterior silk gland, remains intact throughout silk fiber formation and is not replaced by more complex arrangements. Notably, the anisotropic arrangement likely affects the mechanical properties of silk fibers in the x and z directions.

## Discussion and conclusion

In this work, we employed cryo-FIB milling and cryo-ET 3D reconstruction to investigate the native *in situ* hierarchical structure of silk fibers form silkworms, spiders and artificial silk. These findings provide new insights into the spinning mechanisms and the relationship between ultrastructure and mechanical properties, which offer potential pathways for the manufacture of high-performance artificial fibers. Our cryo-ET data conclusively demonstrate that the fundamental structural unit of silk is a nanofibril approximately 3.6 nm in diameter. These nanofibrils are anisotropically arranged, forming a herringbone pattern rather than being merely stacked in parallel. Multiple layers of this herringbone arrangement align in the same direction, ultimately giving rise to micro-scale silk fibers. While spider silk shares this parallel nanofibril organization, it features a notably denser structure compared to silkworm silk (Fig. 5).

**Figure 5.**
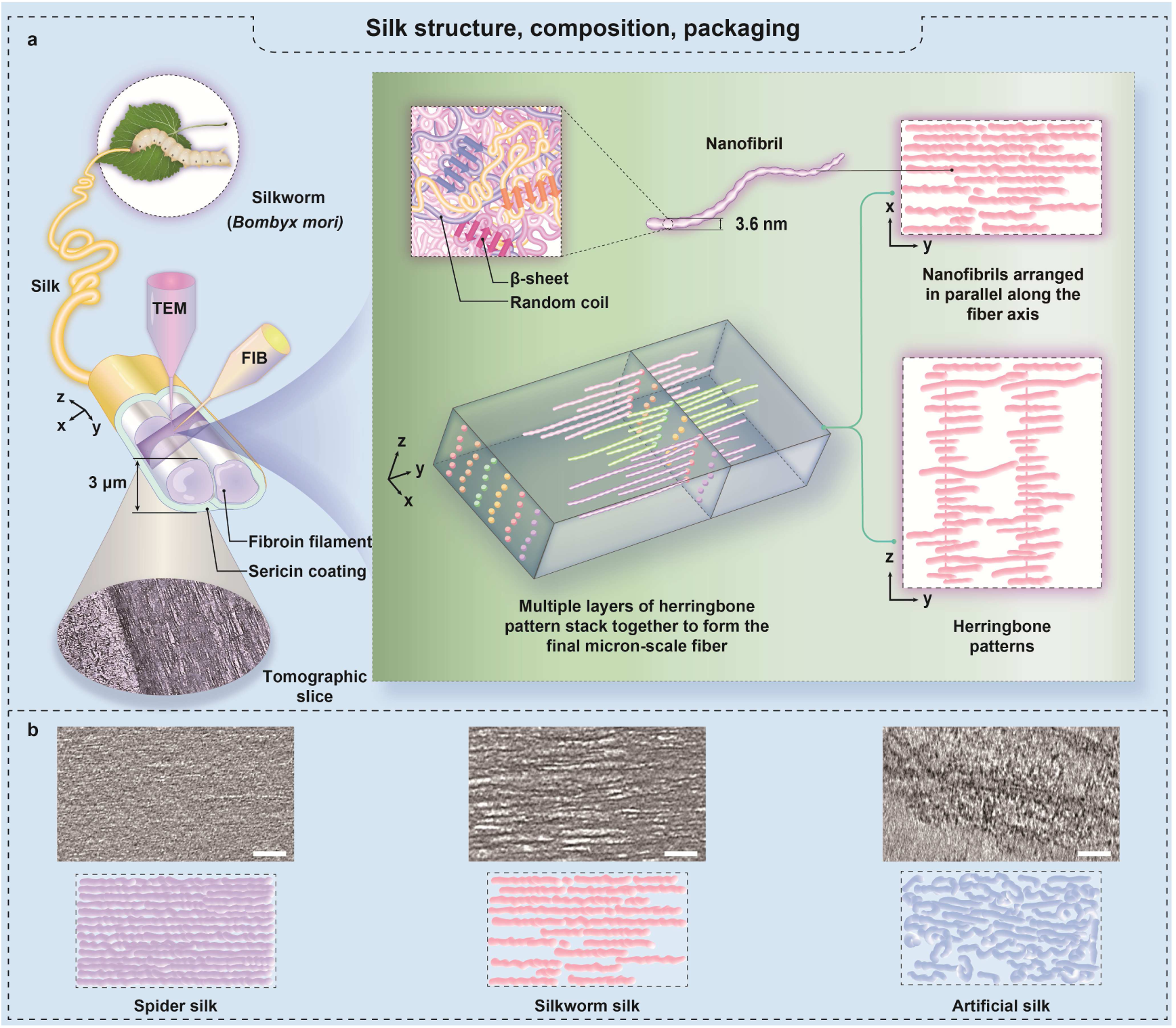
Structure, composition and organization of silk. **a**. The fundamental structural unit of silk is a nanofibril approximately 3.6 nm in diameter. The fibroin is a fibrous protein rather than globular protein, while sericin does not exhibit fibrous characteristics. These nanofibrils within the silk are anisotropically arranged, forming a herringbone pattern rather than being merely stacked in parallel. Multiple layers of this herringbone arrangement align in the same direction, ultimately giving rise to micro-scale silk fibers. **b**. Comparative study of silkworm, spider, and artificial silk. In spider silk, a dense structure featuring nanofibrils is clearly visible, with their long axes tightly aligned parallel to the flow direction, leaving no discernible gaps. By contrast, silkworm silk exhibits larger, regionally varying gaps between nanofibrils, while artificial silk lacks the highly ordered arrangement characteristic of natural silks. Scale bar, 20 nm.

The size, shape, and distribution of fundamental elements are critical to a material’s performance [^64,65^]. There has long been debate about the fundamental elements of animal silk, particularly whether silk proteins are globular or fibrous and what size micelles or nanofibrils form the silk fiber. In this study, we demonstrate that the fundamental unit of silk fiber is a beaded nanofibril with a diameter of approximately 3.6 nm, identified through cryo-ET reconstruction combined with AlphaFold 3 prediction. This is the smallest structural unit currently observable. By avoiding chemical fixation and staining, which risk altering macromolecular organization, we obtained unprecedented detail of silk’s internal structure. However, due to the flexibility of fibroin nanofibrils, we were unable to achieve high-resolution 3D structures through single-particle analysis or subtomogram averaging. Consequently, the exact number of globular structural domains within each nanofibril remains unclear. Nonetheless, it is evident that the more globular domains a nanofibril contains, the greater its flexibility. Therefore, reducing the number of globular domains to decrease the flexibility of nanofibrils, thereby increasing the density of nanofibrils within silk, could be a promising direction for future silk protein gene editing.

Moreover, sericin does not form a regular arrangement during the same spinning process (Fig. 2, a-b), suggesting that the performance of silk fibers is primarily determined by the protein sequence, followed by the spinning process itself. The protein sequence dictates the material’s mechanical properties by shaping the morphology, size and interactions of its fundamental elements. Recent studies have shown that introducing spider silk protein genes into silkworms can produce fibers with enhanced performance, highlighting the significance of the protein sequence [^66^]. Thus, designing protein fibers with specific functional properties through AlphaFold predictions combined with gene editing is another promising direction for future research in silk modification [^14,18,19,67,68^].

The herringbone pattern has increasingly been recognized as a higher-order arrangement [^29,63,69^] with significant implications for the mechanical properties of silk fibers. Whether this pattern is aligned parallel or perpendicular to the fiber axis is crucial for understanding silk fiber assembly. Our study found that the herringbone pattern is perpendicular to the fiber axis, while the nanofibrils within it are arranged parallel to the axis. The exact mechanism of herringbone pattern formation remains unclear. Previous studies suggested that this pattern might be related to interactions between N-terminal domains triggered by pH reduction [^53,63^]. However, our experiments revealed that even in the blank regions of the herringbone pattern, where no nanofibrils were present, a vertical axis perpendicular to the nanofibrils remained (Supplementary Fig. 9a). This suggests that the formation of herringbone patterns may involve some unknown interactions or the participation of other proteins, emphasizing the importance of investigating the transformation of disordered nanofibrils in the silk gland into ordered herringbone patterns for understanding the spinning mechanism and simulating silk fiber formation in vitro. Additionally, we observed variations in the length of the nanofibrils within the herringbone patterns, possibly indicating another type of interaction between the nanofibrils.

Although we were unable to reconstruct the higher-order structures in spider silk due to its dense nanofibril composition, we did observe similar herringbone patterns in the third limb of the spider’s spinning duct, akin to those found in the anterior silk gland of silkworms (Supplementary Fig. 9b). This similarity may indicate a common feature in natural spinning processes. Notably, in both spiders and silkworms, the spinning dope does not completely fill the spinning duct (Supplementary Fig. 9, c-d) [^70^], suggesting a unique liquid environment within these ducts beyond simple acid conditions [^71^]. This specialized environment appears to facilitate the silk extrusion from the spinneret, resembling the process of interface wiredrawing, where continuous fibers are produced by placing protein solutions in suitable buffers. Thus, further investigation into the liquid environment within the anterior silk gland lumen is essential for advancing our understanding of silk spinning mechanisims [^15^].

In summary, cryo-ET of cryo-FIB-milled silks has allowed us to examine the morphology and architecture of silk proteins *in situ*, overcoming artifacts typically introduced during sample preparation with resin, heavy metals, and other agents. These significant advances enabled us to explore the relationship between silk’s ultrastructure and its properties more thoroughly. We unambiguously demonstrated that within silkworm silk, fibroins exist in the form of nanofibrils. These fibroin nanofibrils, with a diameter of approximately 3.6 nm, are the fundamental structural units of silkworm silk, arranged in an anisotropic herringbone pattern. In contrast, the nanofibrils in spider silk are more densely packed. We anticipate that further studies combining cryo-ET of cryo-FIB-milled silkworm silk, artificial silk, and spider silk from seven different spider silk glands, will enable us to better correlate mechanical properties with ultrastructure of the silk, offering new insights into the design and engineering of artificial silk. This study establishes a new research paradigm for the characterization of both natural and synthetic silk fibers, while also providing potential molecular evaluation criteria for assessing the mechanical properties of artificial fibers.

## Materials and methods

### Animals

Silkworms (*Bombyx mori*) were reared to the third larval stage at room temperature, fed a diet of mulberry leaves. Twenty adult female spiders (*Araneus ventricosus*) were collected from Sichuan and Beijing, China, during August and October. The spiders were housed individually in separate containers and provided with moths and water.

### Preparation and vitrification of natural silk fibroin

Silkworm larvae were reared at room temperature until reaching the early wandering stage, after which they were frozen using liquid nitrogen and stored at −80 °C. Silk glands were dissected from the frozen silkworms, and natural silk fibroins (NSFs) were purified from the posterior of middle silk gland (PMSG) using an extraction buffer (50 mmol·L^-1^ NaCl, 50 mmol·L^-1^ Bis-Tris, 0.5 mmol·L^-1^ EDTA, pH 7.5). NSFs were further analyzed by size exclusion chromatography (SEC) using a Superose 6 increase 3.2/300 column (GE Healthcare, USA). The homogeneity of NSFs was evaluated via sodium dodecyl sulfate polyacrylamide gel electrophoresis (SDS-PAGE), followed by Coomassie Brilliant Blue staining. A homemade linear gradient SDS-PAGE gel (3%-16%) was prepared using a Hoefer SG 30 gradient maker (Thermo Fisher Scientific). The concentration of NSF was determined by measuring absorbance at 280 nm (A_280_) with a NanoDrop 2000C spectrophotometer (Thermo Fisher Scientific, USA).

NSF solution was incubated with amphipol (Anatrace, USA) at mass ratio of 1:3, 1:6, 1:9, respectively, at 4°C. Following different incubation periods, the samples were mixed with native-PAGE loading buffer (Invitrogen, USA) and subjected to electrophoresis at 4°C under the following conditions: 90 V for 30 min, followed by 175 V for 90 min. The gels were stained with Coomassie Brilliant Blue. A linear gradient blue native polyacrylamide gel electrophoresis (BN-PAGE) gel (3%-16%) was cast following Witting’s protocol [^72^], with a Hoefer SG 30 gradient maker (Thermo Fisher Scientific).

NSF extracted from PMSG was mixed with amphipol in extraction buffer at a mass ratio of 1:6. After incubation at 4°C for 72 h, 3 μL of the mixture was loaded onto a glow-discharged holey grid (GIG, 1.2/1.3, Au) and vitrified by flash-plunging into liquid ethane using a Vitrobot Mark IV (FEI, USA). Blotting time, force level and humidity were set to be 7 s, 0 and 100%, respectively.

### Metal shadowing

Metal shadowing was conducted as described previously [^29,73^]. For the metal shadowing of spider silk protein, a 5 μL solution (30 μg·mL^-1^) was applied to glow-discharged copper grids for 2 min. Excess solution was carefully removed using blotting paper. For the metal shadowing of NSF *in situ* in the spinning dope of silkworm, a glow-discharged copper grid was placed on the spinning dope and gently pressed with tweezers to avoid any artifact formation. NSF was dehydrated through a graded ethanol series (0, 25%, 50%, 75%, 100%) for 4 min each step. The samples were then shadowed with tungsten using a DV-502B high vacuum evaporator (Denton Vacuum, USA) and air-dried. A tungsten wire (8 cm in length) was clamped between two electrodes at a distance of 3.8 cm, with a separation of 9.3 cm from the center of the specimen platform. The angle between tungsten wire and the sample was approximately 10°. The NSF samples were rotated during evaporation for 14.5 min. Transmission electron microscopy was performed using a Tecnai Spirit (FEI, USA).

### Preparation of silks and Cryo-vitrification

First, 3 μL buffer (50 mmol·L^-1^ Citric acid, 50 mmol·L^-1^ Bis-tris propane, 16% w/v Polyethylene glycol 3,350, pH 5.0) was placed on a glass plate, followed by the addition of an equal volume of NSF solution (3 mg·mL^-1^). The solutions were quickly mixed with a 10 μL pipette tip, and the resulting artificial silks were immediately placed on the glow-discharged mesh grids. Fresh silkworm and spider silks were obtained by forced silking and were also immediately placed on the glow-discharged mesh grids. To prevent dehydration, 3 μL of distilled water was added to the grid before sample application. Excess liquid was blotted with Whatman filter paper for 6 s, and then the grids were rapidly plunge-frozen into liquid ethane using an EM GP system (Leica). Vitrified grids were stored in liquid nitrogen until further used.

### Cryo focused ion beam milling and Cryo-ET data acquisition

Cryo-focused ion beam (cryo-FIB) milling was performed using a Helios NanoLab 600i Dual Beam SEM (Thermo Fisher Scientific), as previously described [^74^]. The goal was to prepare lamellae with a final thickness of ≤ 100 nm. Frozen grids were transferred with the cryo-transfer shuttle into the SEM chamber via a Quorum pp3000T cryo-transfer system (Quorum Technologies, UK) at −180℃. During milling, the angle between the FIB and the silk surface was maintained at 5°-10°, with milling performed from both sides to create vitrified silk lamellae. The ion beam accelerating voltage was set at 30 kV, and the current ranged from 0.43 nA (rough milling) to 10 pA (fine milling). The fine lamellae had a thickness of less than 100 nm.

Cryo-ET data acquisition was carried out using a Titan Krios transmission electron microscope fitted with a K2 detector. Overview tilt series of silks lamellae were acquired under a magnification of ×105,000, resulted in a physical pixel size of 1.36 Å, under counting mode. Before data collection, the pre-tilt of the sample was determined visually, and the pre-tilt was set to be 10° or −9° to match the pre-determined geometry caused by loading grids. And the tilt range were set to be between −60° to +40° for −10° pre-tilt or −40 to +60° for +10° pre-tilt, with a 3° step, resulting in 33 tilts and 99 electrons per tilt series. The slit width was set to be 20 eV, with the refinement of zero-loss peak after collection of each tilt series, and nominal defocus was set to be −3.5 to −4.5 μm. Fro the fibroin extracted from silk glands, the tilt range were set to be between −55° to +55° with a 3° step, resulting in 35 tilts and 115 electrons per tilt series. All tilt series used in this study were collected using a dose-symmetry strategy-based beam-image-shift facilitated acquisition scheme, by in-house developed scripts within SerialEM software. The rest of the collection details are consistent with those mentioned above.

### Tomogram reconstruction and sub-volume averaging of the fibroin

Motion correction and contrast transfer function (CTF) estimation were carried out in Warp; tilt series alignment was carried out in IMOD and AreTomo. For the fibroin extracted from silk glands, 77 tomograms were binned 4 and apply the deconvolution method in Warp to denoise in order to facilitate the particle picking. To pick filaments, CrYOLO [^75^] was employed to detect the straight filaments in the XY planes of the tomograms. These segments were extracted with a box size of 40 pixels (218 Å) using Warp. The reference was used a featureless cylinder using Dynamo. The sub-volumes were then aligned and classified using RELION 4.0. This averaged structure has an estimated resolution of 24 Å based on the ‘gold-standard’ FSC with 0.143 criterion. The details see supplementary Fig. 2.

For the fibroin *in situ*, due to the low SNR in lamella, the tomograms were reconstructed using SIRT-like filter and binned 2 times implemented in IMOD. And the particles were picked by CrYOLO and extracted from the filter tomograms. These segments were extracted with a box size of 90 pixels (245 Å) using RELION. The reference was used a featureless cylinder using Dynamo. The sub-volumes were then aligned and classified using RELION 3.0. This averaged structure has an estimated resolution of 22 Å based on the ‘gold-standard’ FSC with 0.143 criterion. The details see supplementary Fig. 5a.

### Restoring the tomograms by deep learning

Due to the dense arrangement of silk fibers and the presence of a significant missing wedge effect 3D especially in XZ plane, we employed the REST method [^62^] for tomogram restoration. First, we generated a ground truth by randomly distributing the averaged map (many fibers) to densely arrange in the particle box. Then employing andom rotations and shifts (high-quantity datasets), noise and missing wedge artifacts were then added to the ground truth to form the low-quantity datasets. These data were paired and then were used as training pairs for REST training. The trained model was subsequently utilized to recover the raw tomograms from the weighted back-projection (WBP) reconstruction. The details see supplementary Fig. 5b.

### Measurement and Visualization

Visualization and figure generation were performed in UCSF ChimeraX v1.3 and IMOD. The diameter and length of fibroin nanofibrils were measured manually using the measure tool in IMOD and Fiji. To analyze the density of spider and silk fibers, we projected the same number of XY slices into a single 2D projection keep the same volume. The resulting projection was then binarized, and the proportion of the fibers in space was quantified by calculating the ratio of white pixels (representing fibers) to the total number of pixels in the projection.

## Acknowledgments

This work was supported by grands from the National Natural Science foundation of China (32302816, 32241029), the Chinese Ministry of Science and Technology (2021YFA1300100, 2023YFA0913400), and the Chinese Academy of Sciences (CAS) (XDB3700000, JZHKYPT-2021-05). All EM data were collected and processed at the Centre for Bio-imaging (CBI), Institute of Biophysics (IBP), Chinese Academy of Sciences (CAS). We would like to thank Xiaojun Huang, Boling Zhu, for their technical help and support with electron microscopy. Special thanks to Jingwen Song, and Yue Wang, for their help in collecting wild spiders for scientific research.

## Author contributions

K.S., Y.L. and P.Z. conceived the experiments. K.S., Y.L. and X.Z. performed the cryo-EM sample preparation. Y.L. performed the FIB milling. H.Z. and Y.L. performed data collection. H.Z., K.S. and Y.L. analysed the data. K.S. and H.Z. wrote the draft and composed the figures. K.S., H.Z. and Y.L. revised the manuscript and figures. P.Z. supervised the project, analyzed the data and revised the manuscript and figures.

---

**Supplementary Fig. 1.**
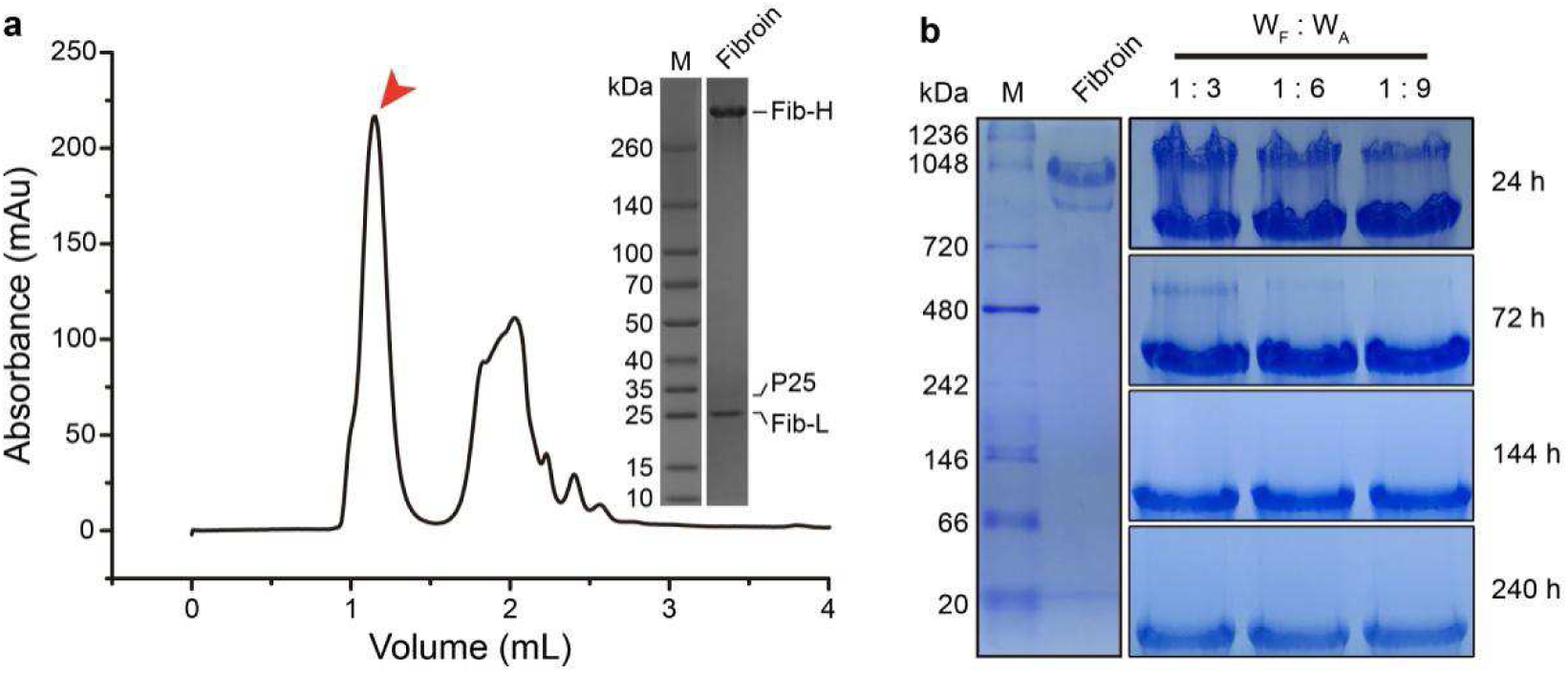
Biochemical analysis of natural silk fibroin. **a**. SEC and 4-16% gradient SDS-PAGE (Coomassie) analysis of natural silk fibroin extracted from the posterior of the middle silk gland, using a Superose 600 Increase 3.2/300 column. **b**. 3-16% BN-PAGE analysis showing changes in natural silk fibroin over time after adding varing mass ratios of amphipol.

**Supplementary Fig. 2.**
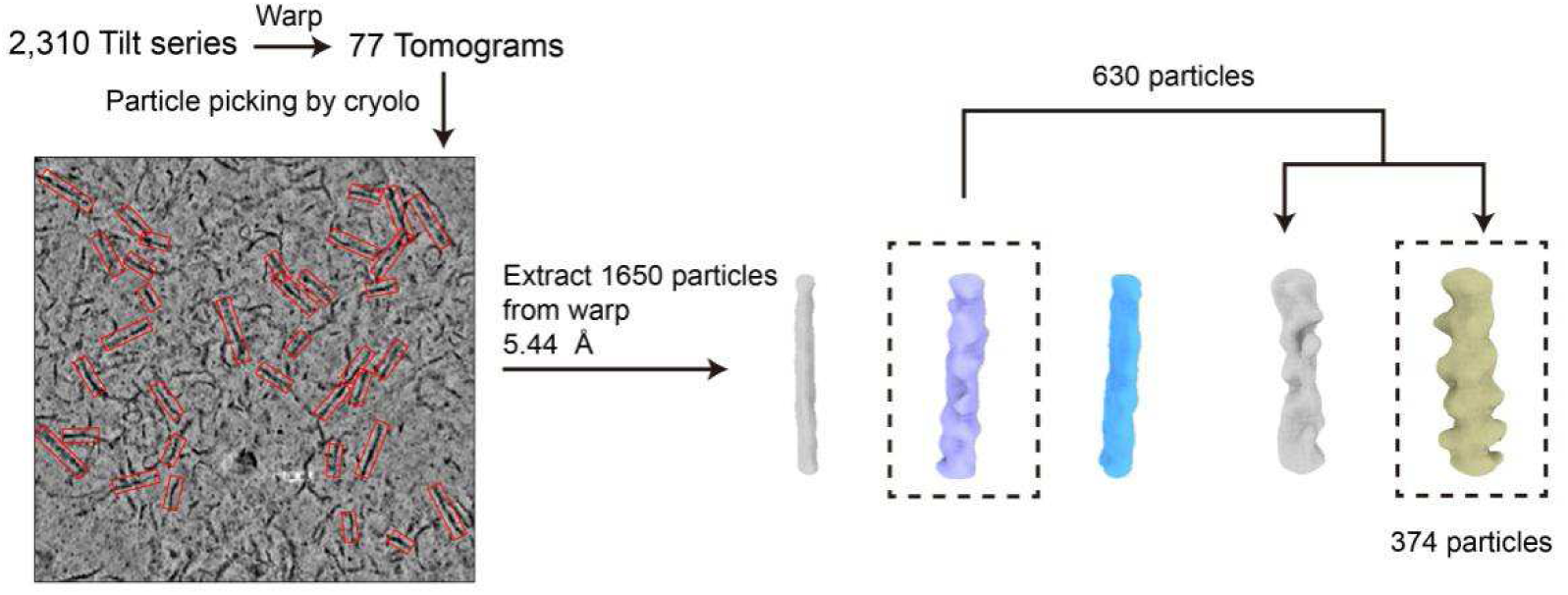
Data processing pipeline, related to Figure 1. Acquisition and processing of natural silk fibroin nanofi brils extracted from the posterior silk glands.

**Supplementary Fig. 3.**
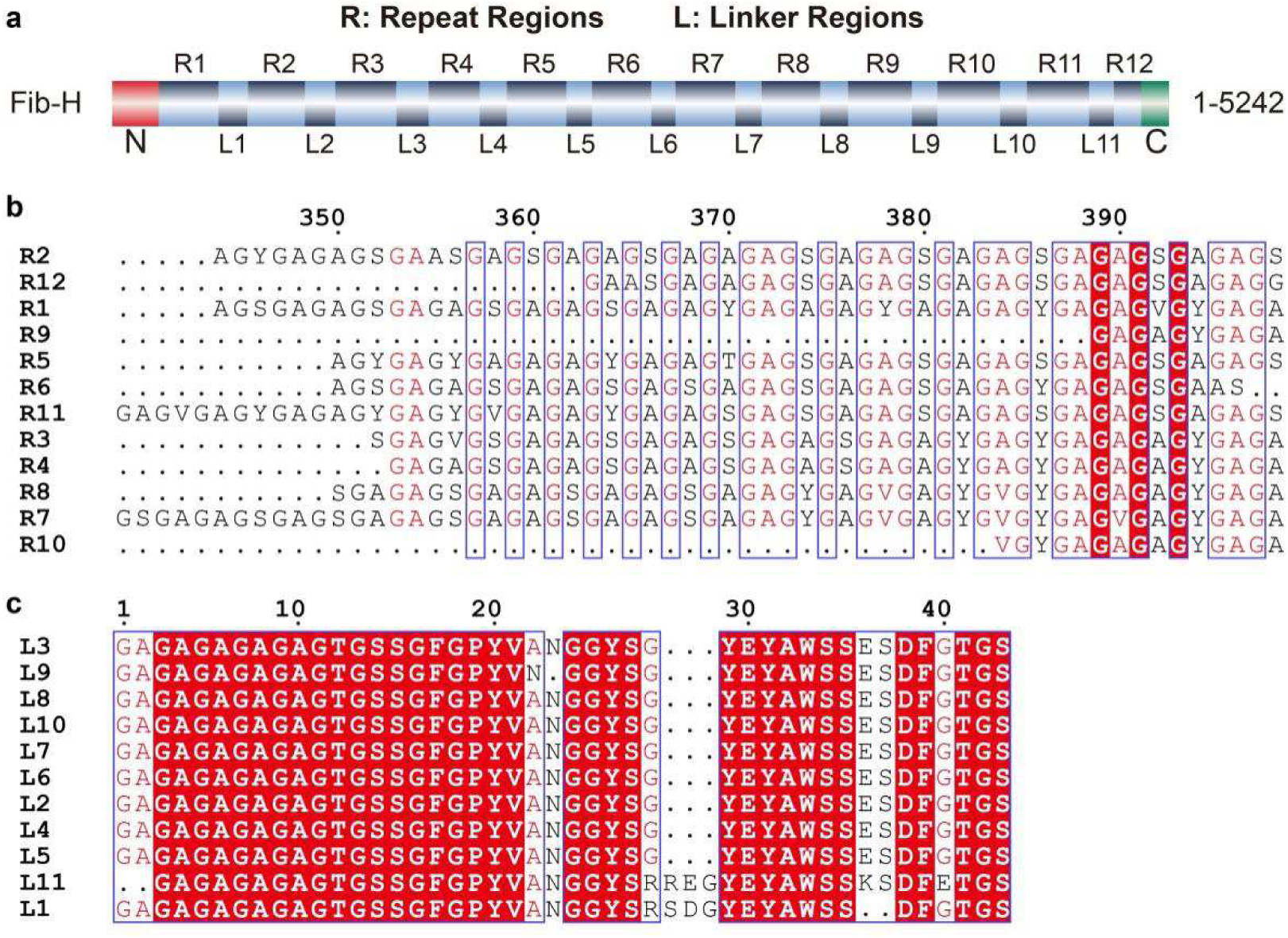
Sequence characteristics of fibroin heavy chain. **a**. Domain map of fibroin heavy chain, illustrating its modular organization with twelve repeat regions linked by eleven linker regions. **b**. Sequence alignment of repeat regions of fibroin heavy chain showing high conservation of the GAGAGS motif. **c**. Sequence alignment of linker regions of fibroin heavy chain showing high conservation of linker sequences.

**Supplementary Fig. 4.**
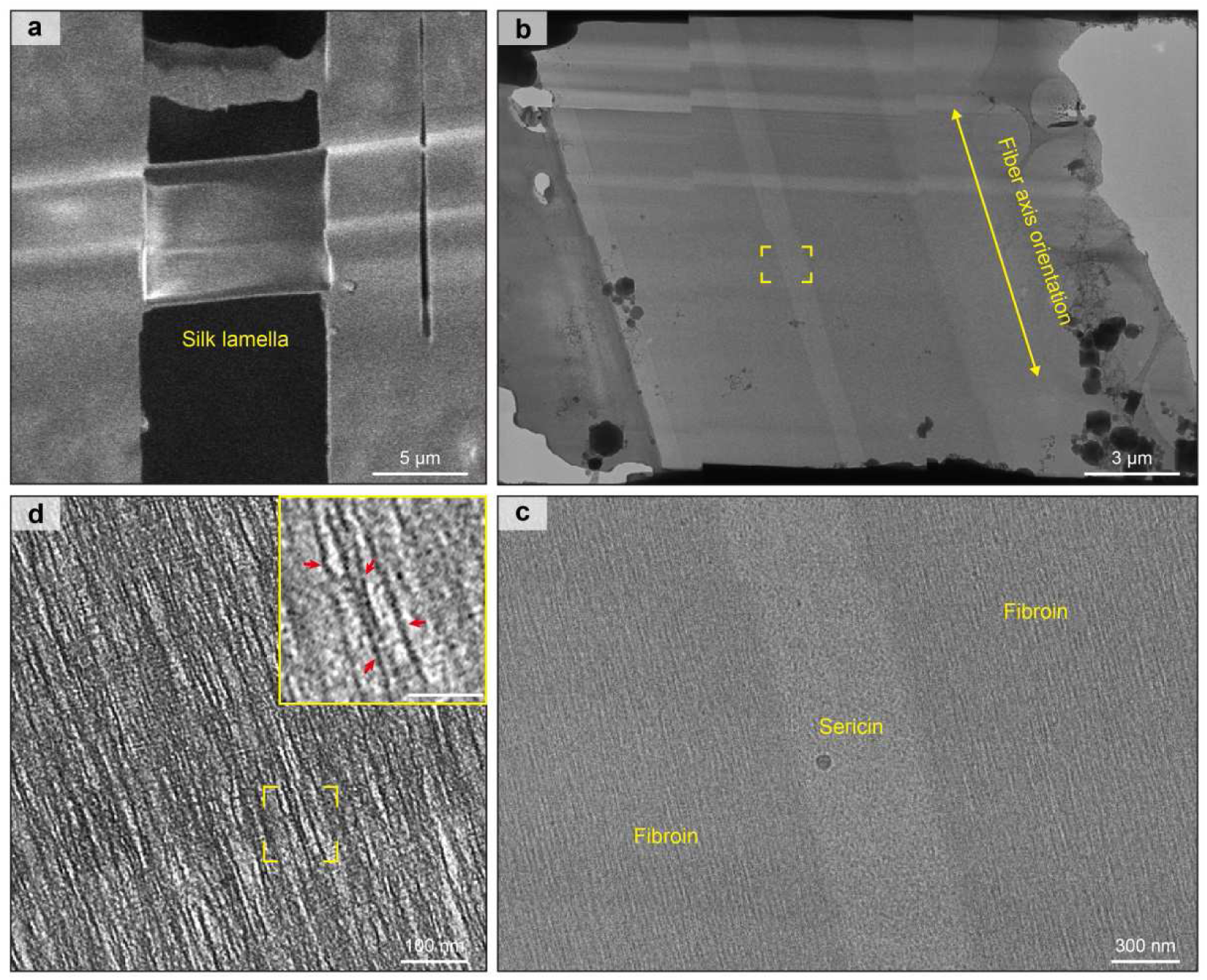
Cryo-FIB-ET of silkworm silk reveals the parallel arrangement of fibroin nanofibrils, related to figure 2 and 4. **a**. Top view of the lamella taken after milling in the FIB-SEM. **b**. The low-magnification projection image of a lamella after FIB milling recorded at the pre-tilt angle of cryo-FIB. **c**. The low-magnification projection image of silkworm silk with representative positioning marked by the yellow box in (b). **d**. Tomographic slice showing parallel arrangement of fibroin nanofibrils, larger view of insets.

**Supplementary Fig. 5.**
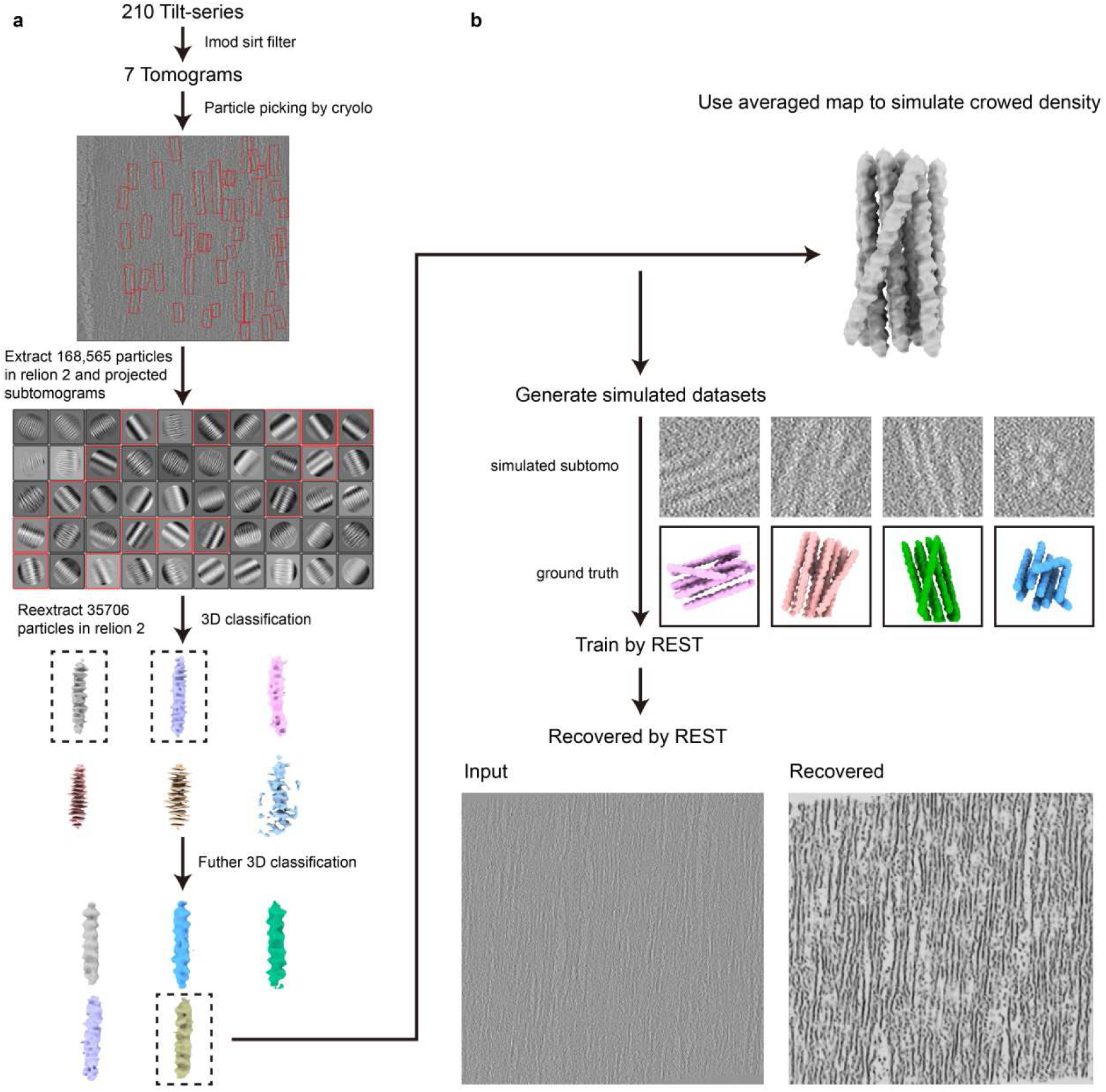
Data processing pipeline, related to Figures 2 and 4. **a**. Processing of fibroin nanofibrils in silkworm silk. **b**. Workflow of REST method.

**Supplementary Fig. 6.**
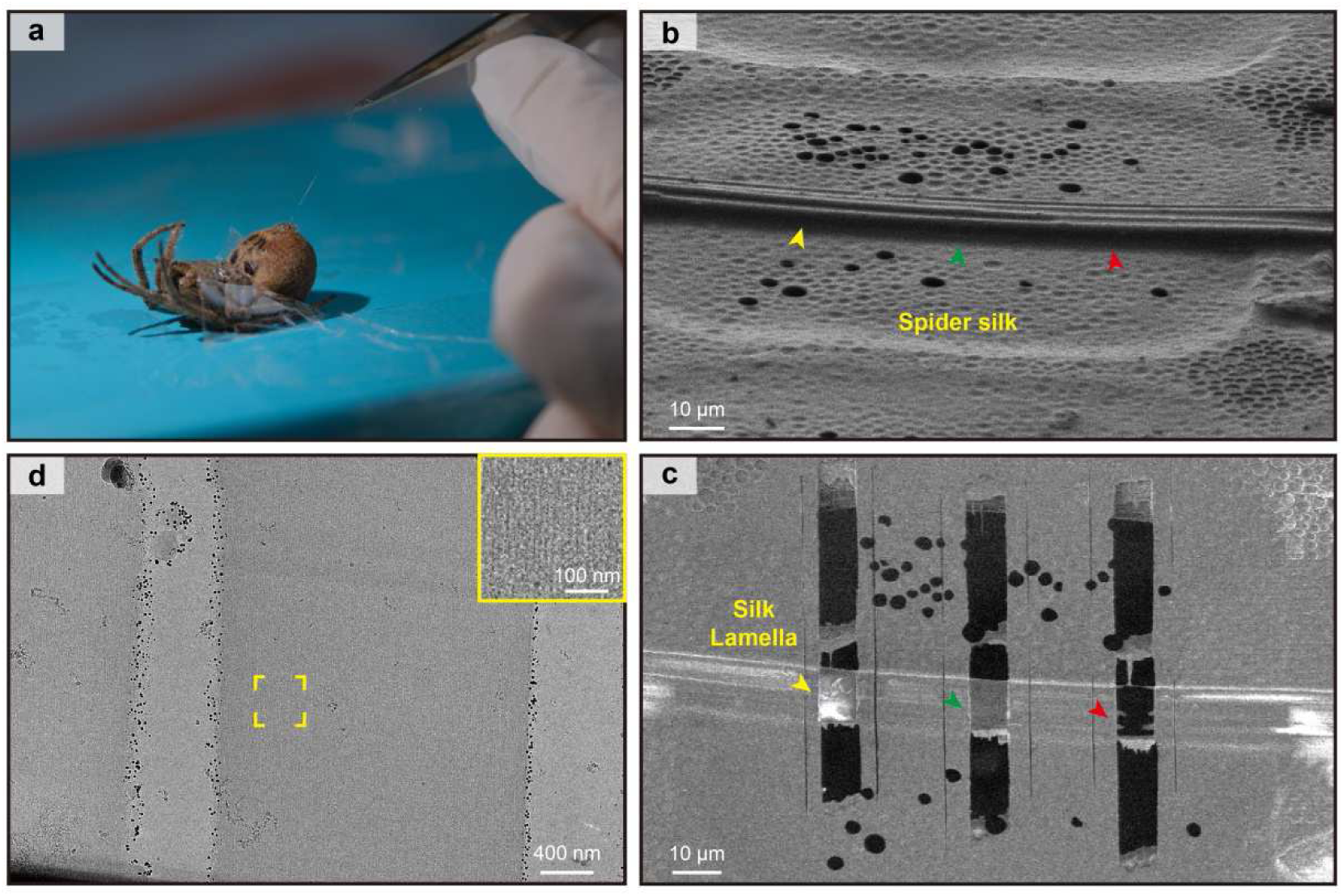
Cryo-FIB-ET of spider silk reveals the parallel arrangement of spidroin nanofibrils. **a**. Silk manually pulled from the spinneret of spider (*Araneus ventricosus*). **b**. SEM image of a grid with vitrified spider silk. The arrowhead indicates the milling position. **c**. Top view of the lamella after FIB-SEM milling. **d**. The low-magnification projection image of a lamella after FIB-milling recorded at a TEM, larger view of insets.

**Supplementary Fig. 7.**
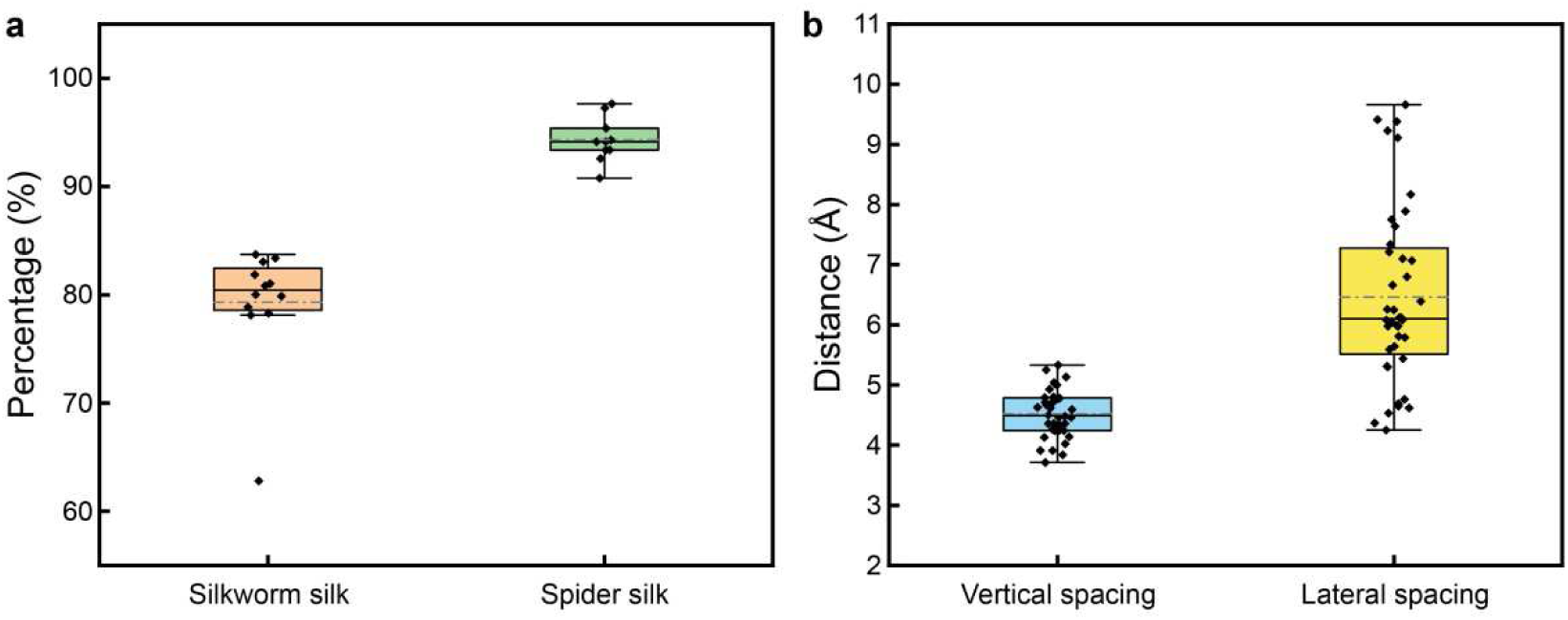
Ultrastructure analysis of silk. **a**. Graph comparing the density of fibroin nanofibrils in silkworm silk and spidroin nanofibrils in spider silk. **b**. Graph shows the distances between adjacent nanofibrils along the z-axis and distances between herriongbone patterns along the x-axis.

**Supplementary Fig. 8.**
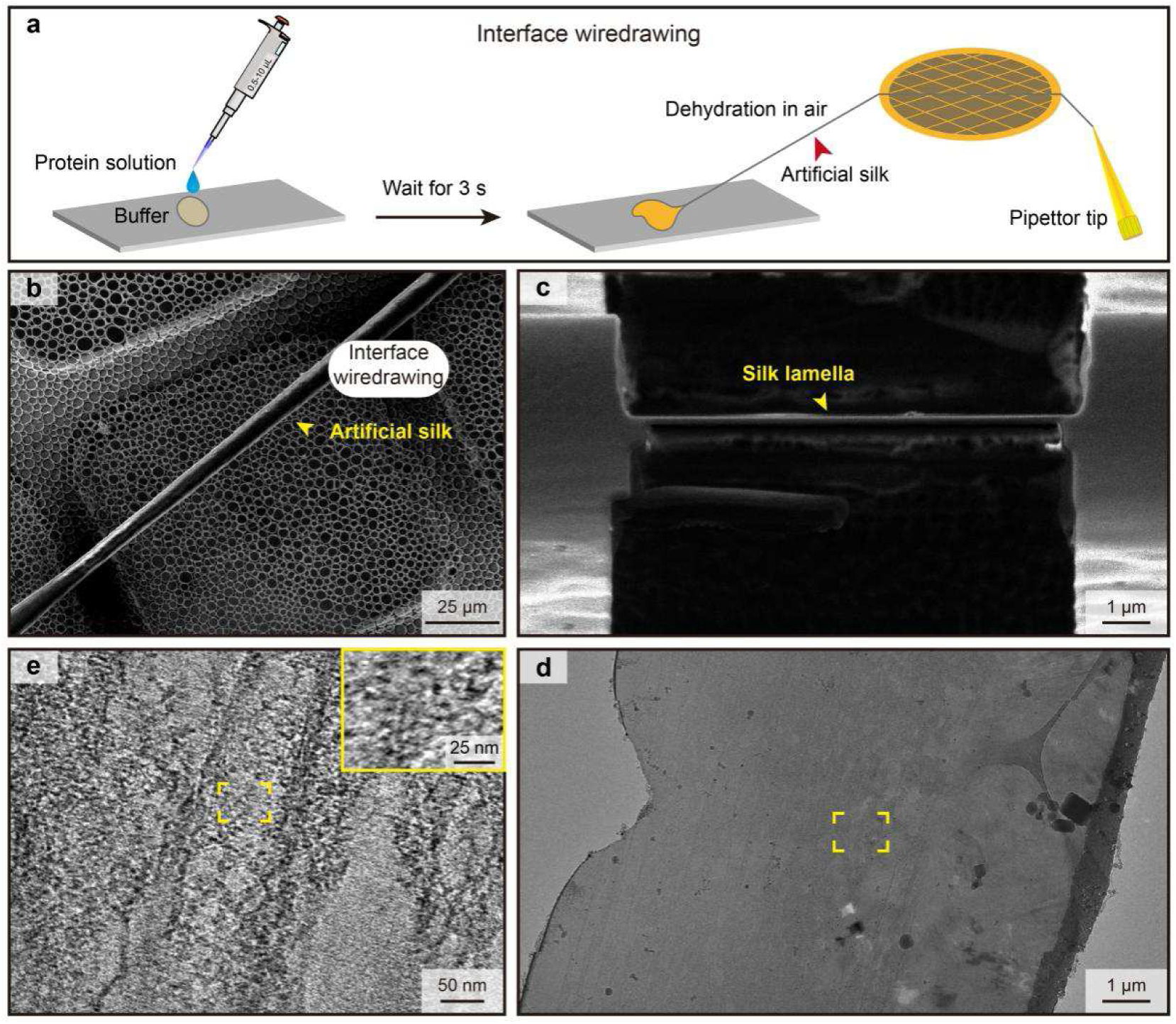
Cryo-FIB-ET of artificial silk reveals a non-uniform phase distribution. **a**. Schematic presentation of artificial silk preparation. **b**. SEM images of the artificial silk. **c**. Side view of the lamella after FIB-SEM milling. **d**. Projection image of a lamella after FIB-milling recorded at a TEM. **e**. Slice through a tomogram depicting a non-uniform phase distribution.

**Supplementary Fig. 9.**
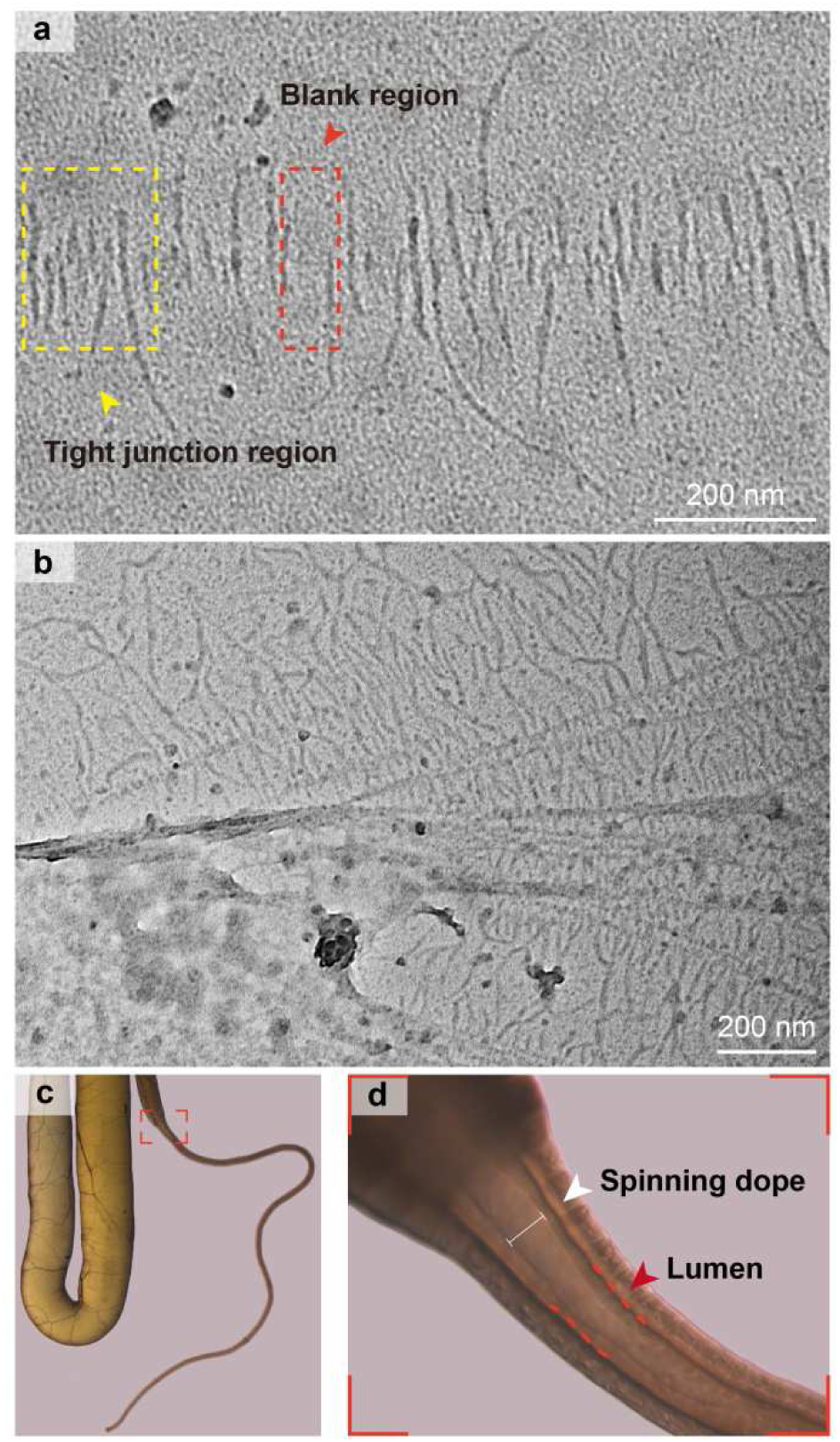
Herringbone patterns in silk glands. **a**. Herringbone patterns formed at the distal of the anterior silk gland of silkworms; the yellow box indicates the tight junction region, and the red box indicates the blank region. **b**. Herringbone patterns formed at the third limb of the spider spinning duct. **c**. Silkworm silk gland; the red box marks the junction between the anterior and middle silk glands. d. Larger view of insets described in (c).

